# Synchronized genetic activities in Alzheimer’s brains revealed by heterogeneity-capturing network analysis

**DOI:** 10.1101/2020.01.28.923730

**Authors:** Sharlee Climer, Alan R. Templeton, Michael Garvin, Daniel Jacobson, Matthew Lane, Scott Hulver, Brittany Scheid, Zheng Chen, Carlos Cruchaga, Weixiong Zhang

## Abstract

It is becoming increasingly evident that the efficacy of single-gene computational analyses for complex traits is nearly exhausted and future advances hinge on unraveling the intricate combinatorial interactions among multiple genes. However, the discovery of modules of genes working in concert to manifest a complex trait has been crippled by combinatorial complexity, genetic heterogeneity, and validation biases. We introduce Maestro, a novel network approach that employs a multifaceted correlation measure, which captures heterogeneity, and a rigorous validation method. Maestro’s utilization for Alzheimer’s disease (AD) reveals an expression pattern that has virtually zero probability of simultaneous expression by an individual, assuming independence. Yet this pattern is exhibited by 19.0% of AD cases and 7.3% of controls, establishing an unprecedented pattern of synchronized genetic activities in the human brain. This pattern is significantly associated with AD, with an odds ratio of 3.0. This study substantiates Maestro’s power for discovery of orchestrated genetic activities underlying complex traits. More generally, Maestro can be applied in diverse domains in which heterogeneity exists.

**Highlights:**

- Synchronized genetic activities associated with Alzheimer’s disease
- Novel vector-based correlation measure that captures genetic heterogeneity
- Enhanced network model for revealing combinatorial genetic interactions
- Pro-survival genetic activities associated with Alzheimer’s disease
- General approach for revealing patterns in data subject to heterogeneity

## Introduction

The manifestation of complex traits involves the intricate interactions of many genes, thereby impeding scientific inquiry. Direct identification of groups of interacting genes that are associated with a trait amounts to examination of an exponential number of gene combinations, which is computationally prohibitive. A general approach to this problem is to build networks of genes so that relationships among the genes can be inferred to aid the identification of interacting or associated genes. However, construction of regulatory networks using methods such as Bayesian networks^1^, linear models^2^, and Boolean networks^3^ requires large amounts of training data and must obey some statistical and/or network restrictions^4^. On the other hand, co-expression network analyses do not impose these requirements. In an early pioneering work of Eisen et al.^5^, Pearson’s Correlation Coefficient (PCC) was used to quantify the correlation between the expression patterns for each pair of genes, thereby identifying pairs whose expression states did not appear to be independent of each other. Based on the PCC values, the genes were grouped using hierarchical clustering, and subsequently mapped into a linear ordering that was visually examined for clusters of co-expressed genes^5^. This method hinges on the well-known notion of guilt-by-association^6^. A key underlying assumption in this network-based approach is transitivity, such that if gene *A* is correlated with gene *B*, and gene *B* is correlated with gene *C*, then *A* and *C* are assumed to be correlated.

It is common for many genes to exhibit differential expression (DE)^7^ between cases and controls when examined in isolation. A typical gene expression “signature” is comprised of a subset of DE genes. While some signatures are merely a collection of DE genes, sophisticated co-expression network methods have evolved over recent years to refine these results and group together potential functionally interacting genes^4,8–10^. In a co-expression network, a node represents a DE gene, an edge represents a significant correlation between expressions of a pair of DE genes, and a group of highly connected nodes captures possible functional associations among the genes. The initial construction of the network typically produces a connected network or graph. Then a clustering method or network community identification method is applied to partition this component into subsets of highly connected nodes, with the goal of identifying *modules*, or *communities*, each of which is a densely connected group of nodes with minimal inter-modular connectivity^11^.

While co-expression network methods are designed to deal with a multiplicity of genetic factors, they ignore genetic heterogeneity, in which different subsets of individuals develop the same trait due to distinct genetic variations and activities. Genetic heterogeneity can cripple the current co-expression network and other network-based analyses as they use conventional correlation measures to determine edge placement^10^. PCC and other correlation measures, such as Euclidian distance, dot product, and mutual information, compute a single scalar. Scalars are insufficient for capturing true correlations in the presence of genetic heterogeneity. If two genes are highly correlated for a subset of individuals but not at all correlated for the others, the correlation value is reduced due to the latter individuals (Figure 1).

**Figure 1.**
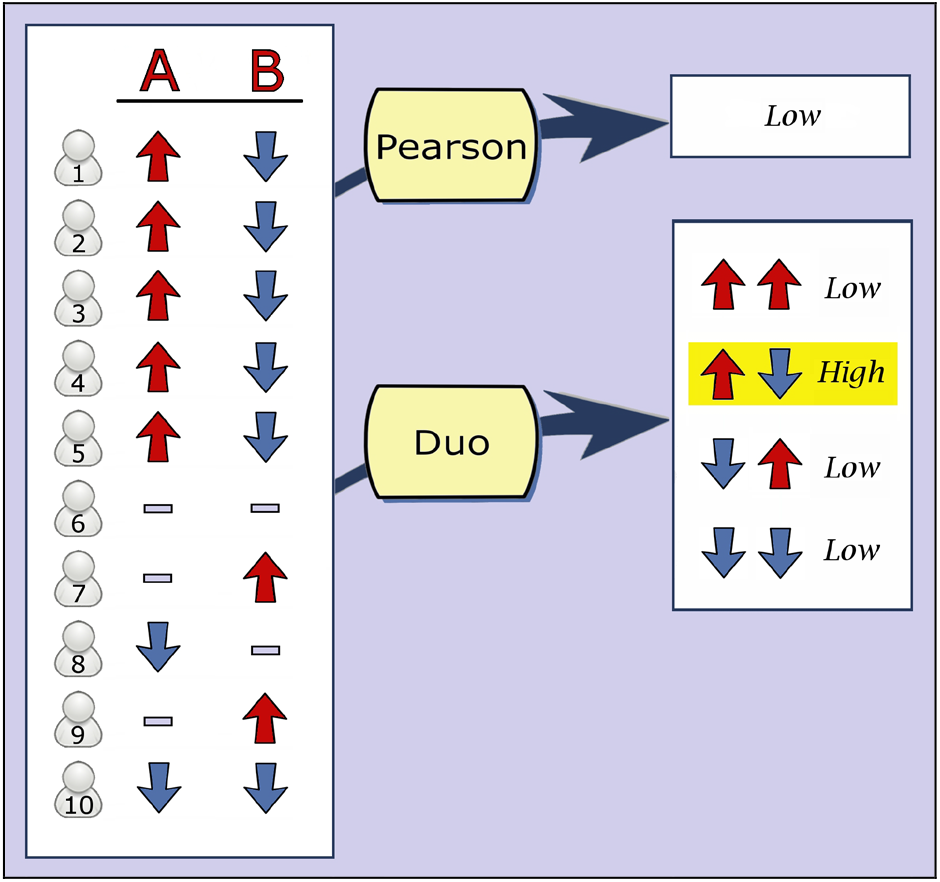
Expression levels and correlation values for genes *A* and *B* for ten individuals. A dash indicates neutral expression. Despite the perfect correlation of the first five individuals, the absolute value of Pearson’s Correlation Coefficient is 0.44 due to the uncorrelated individuals. Duo returns a vector of four values, with the correlation value corresponding to high expression of *A* and low expression of *B* having a highly significant score of 0.80.

We present Maestro, which addresses genetic heterogeneity, exposes combinatorial co-expressions in a decisive and unbiased manner, and rigorously validates the results. Our method diverges from conventional approaches in several ways. First, it hinges upon a novel correlation measure, *Duo*, which we introduce to address an abstruse issue that the conventional methods fail to address, i.e., what constitutes correlation in the face of heterogeneity? To address this issue, our correlation measure computes an autonomous, multifaceted correlation, in the form of a vector, which respects the heterogeneity of distinctive subsets of individuals. An example Duo vector is shown in Figure 1.

Second, Maestro maximizes information retention by substantially enlarging network scaffolding. This expansion provides detailed information about the specific configurations of expression patterns in a highly efficient manner and also reduces false-positive signals such as those due to *dual-expressed nodes* (Figure 2). Dual-expressed nodes arise in conventional co-expression networks as each gene is represented by a single node. This is an insidious flaw as high expression of a given gene may be part of a healthy biological pathway, while low expression might contribute to a deleterious pathway involving a different set of genes, and conventional co-expression networks combine both sets of genes together in a single module due to dual-expressed nodes involved. Maestro eliminates the errors introduced by dual-expressed nodes as it creates two nodes for each gene, each of which represents an expression direction (Figure 2b).

**Figure 2.**
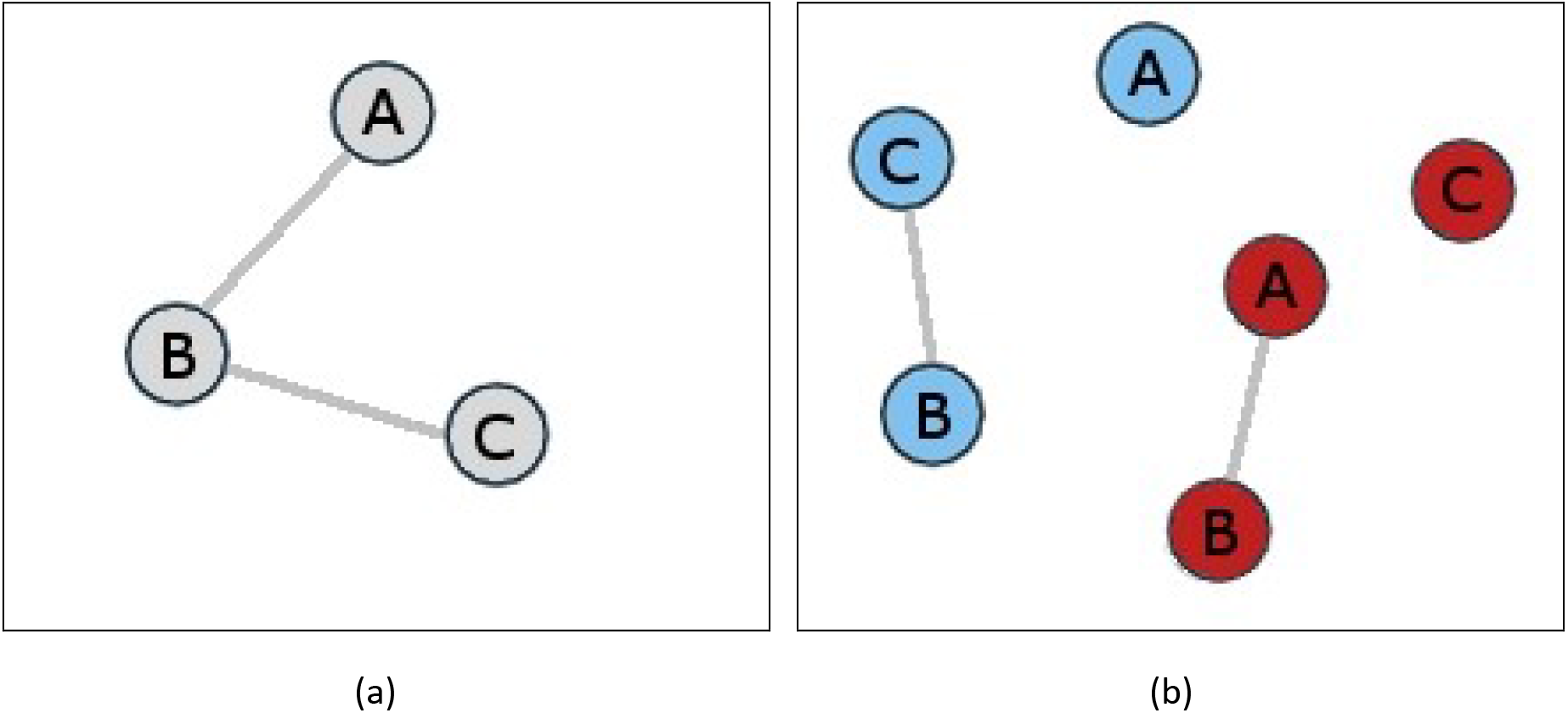
Dual-expressed nodes. This figure depicts three genes, *A*, *B* and *C*, where high expression of gene *A* is correlated with high expression of gene *B* for one subset of individuals, and low expression of gene *B* is correlated with low expression of gene *C* for a different subset. (a) A conventional co-expression network has *A* and *C* connected, via *B*, whereas such a connection cannot be properly established since *B* is involved in two different types of correlations. (b) Maestro networks represent each gene by two nodes in the network, representing high and low expression, respectively (high/low expression is shown with red/blue colors in the figure). This enlargement of the network scaffold removes false positive signals of the type depicted in (a), in which genes *A* and *C* are incorrectly linked by transitivity.

Third, Maestro identifies co-expression modules from the entire sample, including both cases and controls, and then tests the associations of these modules with trait status. In contrast, standard approaches start by testing each gene for DE between cases and controls and thereby produce networks given the trait status. Diagnostic tools and risk prediction are more closely related to the trait associations given the patterns (the Maestro approach) than upon the patterns given trait status (the existing approaches). Indeed, by testing for significant single locus marginal differences in expression between cases and controls prior to building the network, it could be argued that the traditional approach does not truly look at co-expression and instead introduces artifacts by increasing inter-correlations due to shared differential marginal expressions. Furthermore, by examining each gene in isolation with respect to the rest, it is prone to overlook genes with low marginal effects and fail to detect combinatorial effects of a group of genes, defeating the ultimate goal of co-expression network analysis. In contrast, Maestro identifies combinatorial expression patterns without regard for trait status. Hence, Maestro addresses more epidemiologically-relevant questions than traditional approaches and can generate viable biological hypotheses by a small number of trials between cases and controls, which are designed to gain statistical power and enhance discovery of associations of genetic variations and traits.

Fourth, it should be noted that the Maestro networks we have constructed are comprised of distinct modules that do not require the selections of clustering methods and associated parameters.

Finally, Maestro performs multiple rigorous and unbiased validation trials, thereby providing solid information worthy of further research efforts. We have applied Maestro to late-onset Alzheimer’s disease (AD) gene expression data and discovered an unprecedented large-scale pattern of synchronized genetic activities in the human brain that exhibits highly significant association with AD.

## Results

We examined gene-expression profiling data from human cortex tissue for 8,560 genes for 176 AD cases and 188 controls^12^. We defined the highest and lowest 25% of expression values in the total sample as “high” and “low” expression, respectively. The middle 50% of values were regarded as “neutral.” A single network was built using both cases and controls data, without information linking samples to their group identities. Each gene was represented by two nodes, representing high and low expression, respectively. Correlations for each pair of genes were computed using Duo (see Methods) and edges were placed between node pairs with the 1,000 highest correlation values. (We varied this number to test its sensitivity, see below.) The resultant network was explored using breadth-first search, revealing nine discrete disconnected modules, with no edges connecting them to one another. The numbers of individuals exhibiting expression patterns corresponding to the modules were tallied and the odds ratio and Bonferroni-corrected p-value based on the G-test were computed (see Methods).

### Network structure

The 1,000 highest Duo correlations fell into nine discrete modules (Figure 3). The overall network is highly sparse as 16,836 of the 17,120 nodes (98.3%) were *singletons* without adjacent edges. Five of the nine modules were *doubletons*, each of which is comprised of two nodes connected by a single edge. In general, the modules were surprisingly dense, despite the overall sparseness of the network. The largest module had 136 nodes with 583 edges and the second largest had 116 nodes with 371 edges. Supplementary Table S1 has a complete description of this network. Notably, these results demonstrated a strong community structure that arose naturally, without the aid of clustering methods, as we observed distinct dense modules amidst a vast number of singleton nodes.

**Figure 3.**
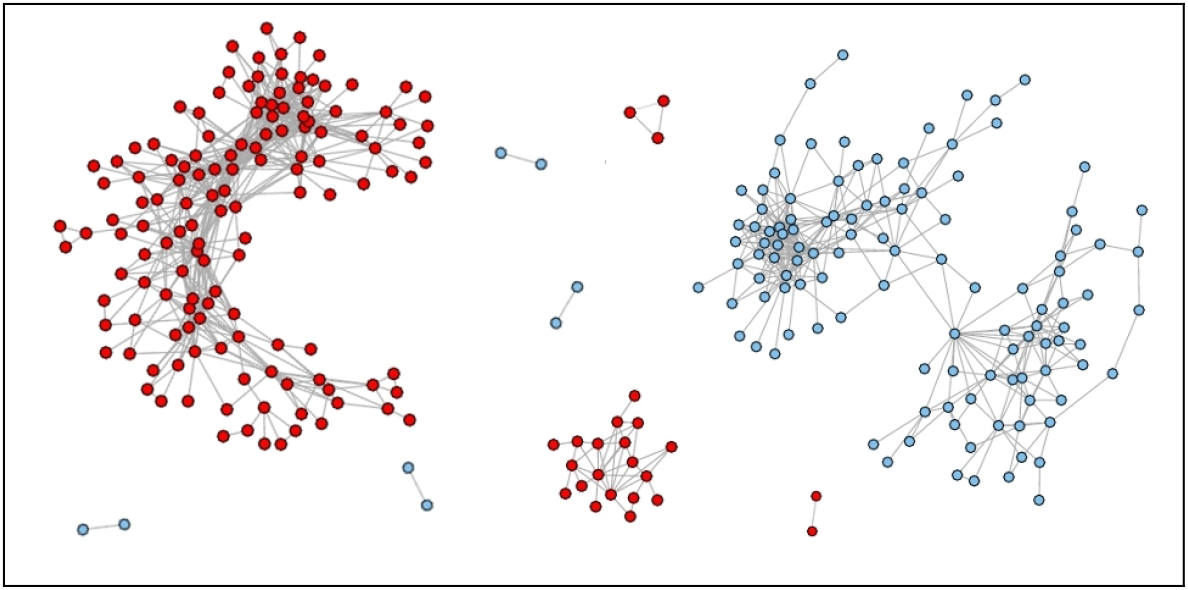
Maestro network consisting of nine communities of genes. Shown is the network built from 364 samples (176 AD cases and 188 controls). Red and blue nodes represent high and low expression, respectively. Each edge represents a significant Duo correlation between the two corresponding nodes; the edges corresponding to the 1,000 highest Duo values are shown. The entire network consisted of 17,120 nodes, among which 16,836 were singletons with no incident edges and are not shown for clarity. The largest community is comprised of 136 genes with high expression, while the second largest consists of 116 genes with low expression. These two modules have 107 genes in common, only the expression direction is reversed.

### Gene expression patterns

Each of the modules in the network defines a gene expression pattern inferred from the entire sample of all individuals. We tested whether any of these patterns were actually exhibited by any of the individuals in the sample and compared frequencies of these combinatorial expressions between cases and controls using a *Carriers* algorithm (see Methods). Five of the nine modules, including the two largest modules, had odds ratios of at least 1.5 and corrected p-values no greater than 3.9×10^−39^ (see Supplementary Spreadsheet S1).

All of the 136 genes in the largest module had high expression. Supplementary Spreadsheet S1 lists these genes and the frequencies of high expression for cases and controls. Given these frequencies and assuming independence, the probability that an individual would possess this entire pattern is 1.07×10^−48^ for cases and 1.34×10^−81^ for controls. Despite these remote odds, 19.0% of the cases and 7.3% of the controls possess this entire expression pattern. These results suggest these genes are not expressed independently of each other. The appearance of a synchronized pattern of this size for so many individuals is unprecedented, and the significant variation between AD cases and controls is noteworthy, yielding an odds ratio of 3.0 and corrected p-value < 2.3×10^−306^.

We examined potential covariates (gender, age at death, cortical region, APOE status, day of expression hybridization, institute source of sample, postmortem interval, and transcription detection rate) for possible associations with this pattern. Supplementary Spreadsheet S2 contains the states of these variables for the cases and controls, with individuals possessing the 136-gene expression pattern highlighted. Only one of these properties exhibited any association with possession of the 136-gene pattern -- the APOE genotype. We found a significant effect on those with the *ε*4 allele; namely, if a subject has the *ε*4 allele, they are more likely to exhibit the 136 pattern if they are a case than if they are a control (p = 0.0266), but there is no significant effect if an individual does not have the *ε*4 allele (p = 0.0672), under the standard significance cutoff of 0.05. Consequently, there is an APOE genotype effect, but it is not strong enough to explain the differences in the frequency of 136-gene expression pattern in cases versus controls.

The second largest module in the network includes 116 genes, had an odds ratio of 1.5, and corrected p-value < 2.3×10^−306^. This pattern exhibits a protective association as it appears significantly more often in controls than AD cases (note that we use the inverted formulation of odds ratio, as described in the SI). Similar to the largest module, this pattern is remarkable as the probability that an individual would possess the entire expression pattern is 1.81×10^−70^ for cases and 1.70×10^−45^ for controls if independence is assumed. Despite these low probabilities, the entire pattern is possessed by 8.7% of the cases and 12.4% of the controls. Of particular interest is the fact that 107 of the 116 nodes in this module appear in the largest module but with opposite expression direction. In summary, the two large modules in the network represent risk and protective patterns and share most of their genes, but with opposite expression directions. The natural appearances of these large, significant modules detected by mining the expression data with no information of sample trait status, cross validate each other.

### Validations

Traditional co-expression network results are typically validated using some type of enrichment analysis based on reference databases such as DAVID^13^ or MetaCore^14^. These methods supply enrichment scores or p-values to indicate the significance of the list of genes. While these scores can be useful, they can also exhibit different types of biases, dependent on which functional annotation resources were utilized and what significance cutoff values were adopted. More critically, because these approaches depend heavily on information of known biological processes and signaling pathways, they may be ineffective or even misleading for poorly understood traits, such as AD. An added complication is the fact that the number and/or sizes of modules may bias the enrichment scores that are calculated^4^. In order to ensure the robustness of our results, we employed a series of rigorous and unbiased trials (see Methods), as follows.

#### Permutation trials

We ran 1,000 permutation trials in which the individuals were randomly sorted for each gene. Consequently, each gene retained the same distribution of values, while inherent correlations were broken up. The largest Duo value across all permutation trials was 0.524 and the minimum Duo value for any of the edges in the original network was 0.740. These results have given a confidence that it is highly unlikely that any of the Duo edges arose by random chance. These results also demonstrated Duo’s ability to discern true correlations from noise.

It should be noted that spurious edges can be introduced due to small sample size, as described in SI. The minimum number of samples needed likely varies for different datasets. For these reasons, permutation tests should be run whenever Maestro is used to ensure that false positive edges are not produced or are as minimal as possible.

#### Replication of results

A confounding issue for most of the published gene expression studies has been the inability to replicate results for independent datasets. Suárez-Fariñas et al. suggested these discrepancies might be due to variations in sample preparation, technician experience, equipment calibration, and choices of pre-processing algorithms, statistical tests, and thresholds^15^. Gudjonsson et al. also noted these types of inconsistencies and suggested they may be due to study design, sample sizes, microarray platforms, and/or batch effects^16^.

Since the original AD dataset was adequately large, we split it in half and used the first half as a “discovery” dataset and the second half for “validation” of the patterns identified by the first half. This approach has the advantage of eliminating some of the technical issues that arise for replication in independent data.

We randomly split the data into discovery and validation sets, each of which had data for 88 cases and 94 controls. We combined the cases and controls data for the first half (discovery data) and built a 1000-edge network using Duo and identified the connected modules using breadth-first search. The network topology was similar to the original results, with eight doubletons, and six modules with three or more nodes, each of which were relatively dense. As in the original results, there were two modules with more than 100 nodes, here they both had 113 nodes. Also, as before, these two modules had opposite regulation patterns and 81 (71.7%) of the genes were the same, just with opposite expression.

The Carriers algorithm was applied to both the discovery and validation datasets using the modules identified in the discovery data network. Supplementary Spreadsheet S3 contains this output and a summary of the two largest modules is summarized in Supplementary Table S2. The largest pattern with high expression was significant as it was possessed by 13.1% and 4.7% of the cases and controls, respectively, in the discovery group. This pattern had an odds ratio of 3.1. This same pattern of 113 genes with high expression was also significant for the replication data, with 20.0% and 7.3% of cases and controls, respectively, and an odds ratio of 3.2.

The pattern of 136 genes with high expression that was found by building the network for the entire dataset of 176 AD cases and 188 controls possessed 104 of the 113 genes identified by using half of the samples in the discovery dataset. Although the 136-gene pattern exhibited a slightly lower odds ratio than the 113-gene pattern, we focus on the larger pattern in this manuscript as it fully utilized all of the data and supplied additional synchronized genes.

#### Variation of network density

We varied the number of edges in the network to test the sensitivities of the modules to our somewhat arbitrary choice of 1,000 edges. For this test, we created four additional networks, in which we reduced the number of edges to 500 and then increased it to 5,000 (5k), 10,000 (10k), and 15,000 (15k). The results are shown in Supplementary Table S3. Note that the networks remained highly sparse as the number of edges in the network increased, and even for the 15k networks, 95.9% of the nodes remained singletons. As more edges were added to the network, the two largest modules tended to capture most of them and for the 15k network, 99.5% of the total edges were located within the two largest modules.

The original 136-node module had 105 nodes in the 500-edge network. For the larger networks, it grew to 212, 242, and finally 268 nodes, with odds ratios greater than 2.4 and p-values < 2.3×10^−306^ throughout. The progression of this module as the networks grew is depicted in Figure 4.

**Figure 4.**
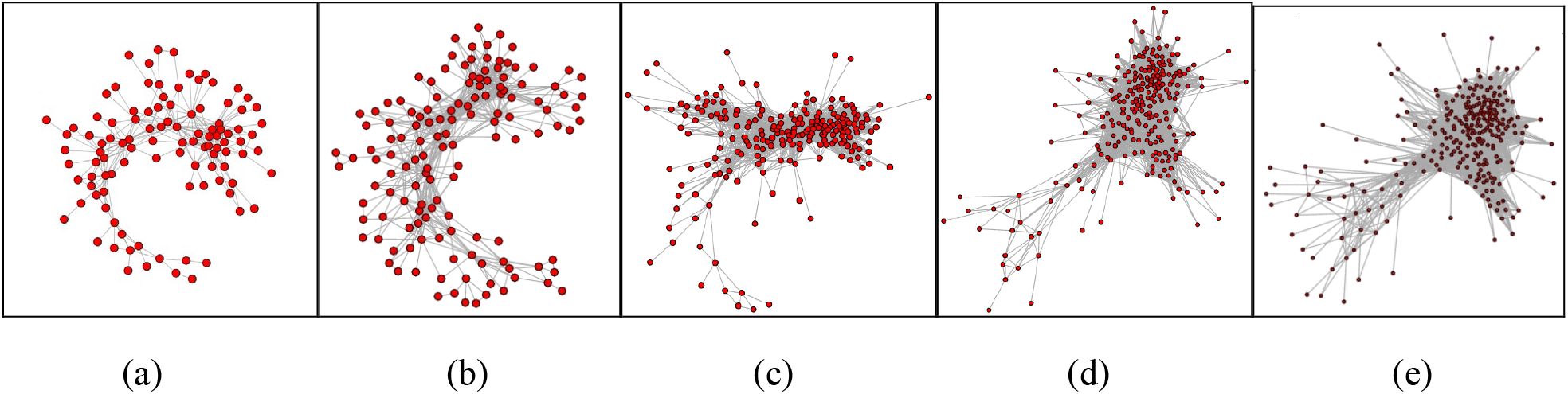
Growth of the largest module with varying network density. The numbers of edges in the network were varied to test the sensitivity of the results to this parameter. Shown are the modules with the most edges for (a) 500-, (b) 1000- (original), (c) 5000-, (d) 10,000-, and (e) 15,000-edge networks. As edges were added to the network, this module grew. However, the number of edges increased considerably more rapidly than the number of nodes. The number of nodes had a fold change of 2.6: from 105 in (a) to 268 in (e); while the edges had a fold change of 24.9: from 319 in (a) to 7,958 edges in (e). For every one of the five networks, more than half of the edges congregated within these modules, reflecting strong combinatorial co-expression regardless of the selected network density.

The original 116-node module had 74 nodes in the 500-edge network, then grew to 185, 220, and finally combined with the third largest module, growing to 341 nodes. For the 10k-edge network, the expression pattern corresponding to this module had an odds ratio of 2.1 and p-value < 2.3e×10^−306^, thereby showing more significant variation than the 116 node module in the original 1k network. This significance reduced after it merged with the third largest module in the 15k-edge network.

There are two criteria to be considered in finding the “optimal” number of edges. First, is the percent of cases affected, and second is the odds ratio. For the largest module, the highest percent of cases is 19%, as found at both 500 and 1k edges, but the 1k module has a much higher odds ratio than the 500. However, the odds ratio does increase when we go to 5k, but at the expense of a substantial drop in the percent of cases. For the second largest module, the highest percent of cases is for the 500-edge network, as well as the second highest odds ratio. The highest odds ratio occurs at 10k, but only with a substantial reduction in the percent of cases. These two criteria can be combined into a weighted odds ratio: the product of the percent of cases times the odds ratio. For example, for the largest module, the weighted odds ratios are (going from 500 to 15k, as in Table S3), 0.456, 0.570. 0.386, 0.291, 0.289. These values indicate that the best results are obtained for the 1k network. For the second largest module, the weighted odds are 0.155, 0.131, 0.083, 0.076, and 0.016, going from 500 to 15k. This indicates the optimal module to use in this case is the one produced by the 500-edge network.

Overall, these trials demonstrated the robustness of the modules when varying the number of edges in the network. As a bonus, they introduced some fine-tuning of the modules that improve upon the case/control variation that was captured in the original network and supply additional synchronized genes.

#### Segregation of data with known heterogeneity

We demonstrated Maestro’s ability to segregate heterogeneity by manually segregating known heterogeneity and testing the subsets. We separated AD cases from controls and built two Maestro networks, one for each group of individuals. The results are summarized in Supplementary Tables S4 and S5 and detailed in Supplementary Spreadsheets S4 and S5. Each network produced a single large module and several small modules with no more than three nodes each. The large modules were highly similar to the two large modules identified in the original network shown in Figure 3, demonstrating Maestro’s ability to segregate heterogeneity.

Specifically, 89% of the genes in the 136-node module from the original network appeared in the 164-node module in the AD cases network. Similarly, 97.4% of the genes in the 116-node module of the original network appeared in the 160-node module that arose in the controls network. These results demonstrate the accuracy of Maestro for segregating heterogeneity as the manual separation of known heterogeneity produced highly similar results to the original network built from all samples combined.

By manually reducing heterogeneity prior to analysis, these results provide additional genes that may be associated with AD. However, unlike the modules that were blindly constructed in the original results, these patterns may be biased. By searching for patterns within a single group, e.g. AD cases, it is not improbable that the identified patterns might be more common for that group. Herein lies the familiar trade-off between reducing possible false positives by increasing false negatives. Maestro is stringent and false positives are highly unlikely, leaving open the potential for false negatives. Nevertheless, we uphold this stringency so that the primary results presented here are worthy of future research efforts.

#### Visual inspection of expression patterns

The carriers algorithm (see Methods) tallies the individuals that possess the entire expression pattern but does not capture individuals that possess most of the pattern. It is possible that partial patterns might be more prominent in one group than the other, potentially weakening the significance of the association. In order to detect these partial patterns, we plotted all of the expression values for the module of 136 genes, as shown in Figure 5. Notice that the risk AD pattern has an “inverse” pattern, shown in blue. This inverse pattern was largely captured by the second largest module in the network (Figure 3). These visual inspections confirm the existence of these large-scale expression patterns and expose their pronounced synchronicity. Moreover, they elucidate the tendency for lockstep expression of these 136 genes, as few individuals possess simultaneous high and low expression of genes in this module.

**Figure 5.**
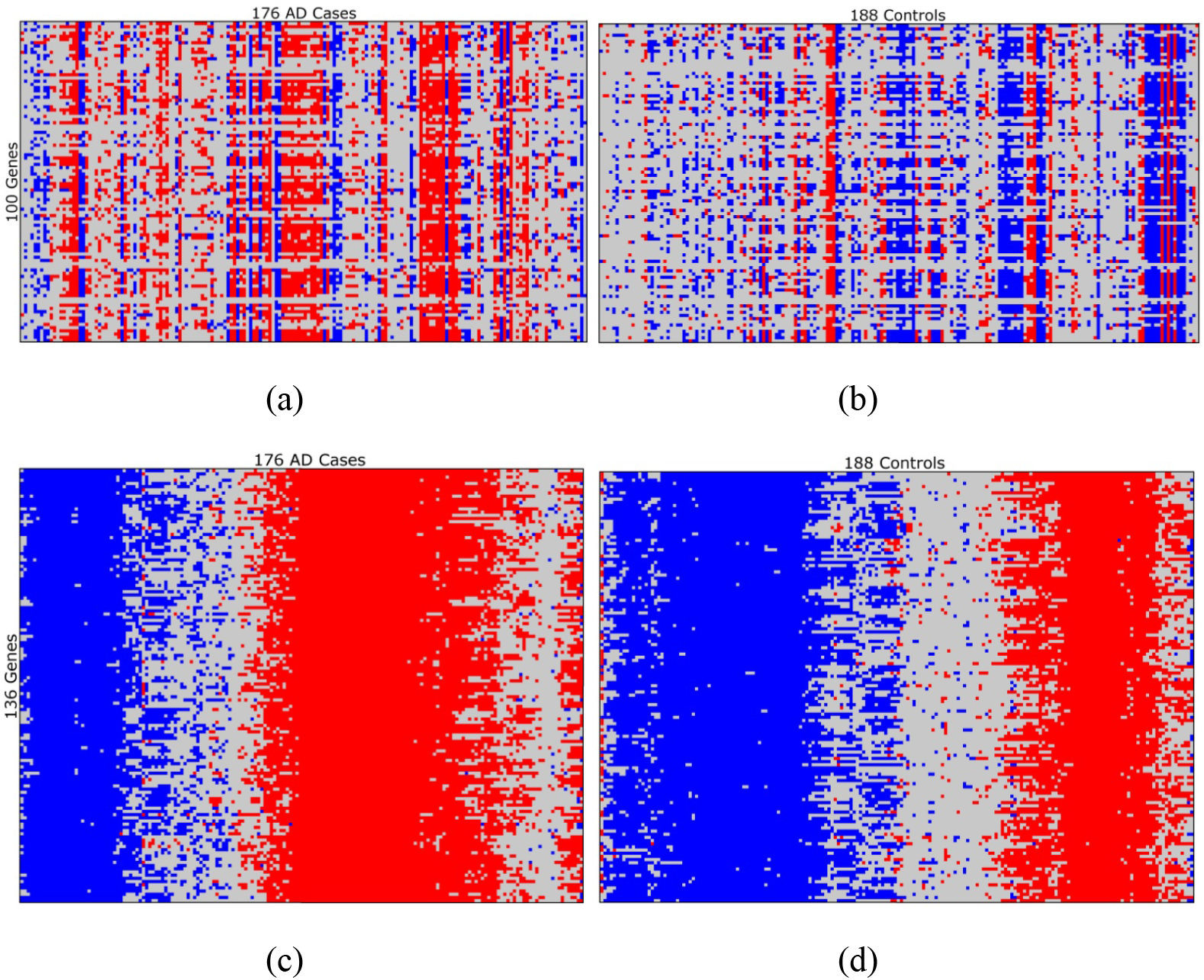
Visual examination of expression levels for selected sets of genes from the AD cases and controls. (Best viewed in color.) Each column represents an individual and each row represents a gene. Red and blue colored cells represent high and low expression, respectively. Light gray indicates neutral expression or missing data. Columns (individuals) have been ordered using TSP+*k* rearrangement clustering^129^ to group similar columns together. Shown is a genetic signature comprised of 100 genes from the AD data for which the cases had the highest differential expression, for the (a) AD cases and (b) controls (identified using SAM^7^). None of the individuals have high expression for all 100 genes, as shown. Maestro was used to identify modules of synchronous genes and shown are the gene expressions in the largest module for the (c) AD cases and (d) controls, which possessed 136 genes. 19% of the cases and 7.3% of the controls had high expression for all 136 genes. Inverse patterns arose spontaneously, as shown in blue. There were a total of 27 missing values in (c) and (d) and each one was in a column that is primarily blue. Due to the fact that a gene with extremely low expression might be labeled as “missing”, increased testing sensitivity might increase the density of the inverse pattern. Note the tendency toward lockstep expression of the genes for the Maestro module, in which most individuals exhibited nearly uniform expression across the 136 genes.

### Comparison with an alternative co-expression network method

We analyzed the data using a well-established co-expression network CoExp method^4,18^ (see SI) and comparisons between Maestro and CoExp are summarized in Figure 6. The resultant AD co-expression network had 1,565 nodes and 15,629 edges. We explore several key differences between the results for the co-expression method and Maestro in SI and summarize these results here.

**Figure 6.**
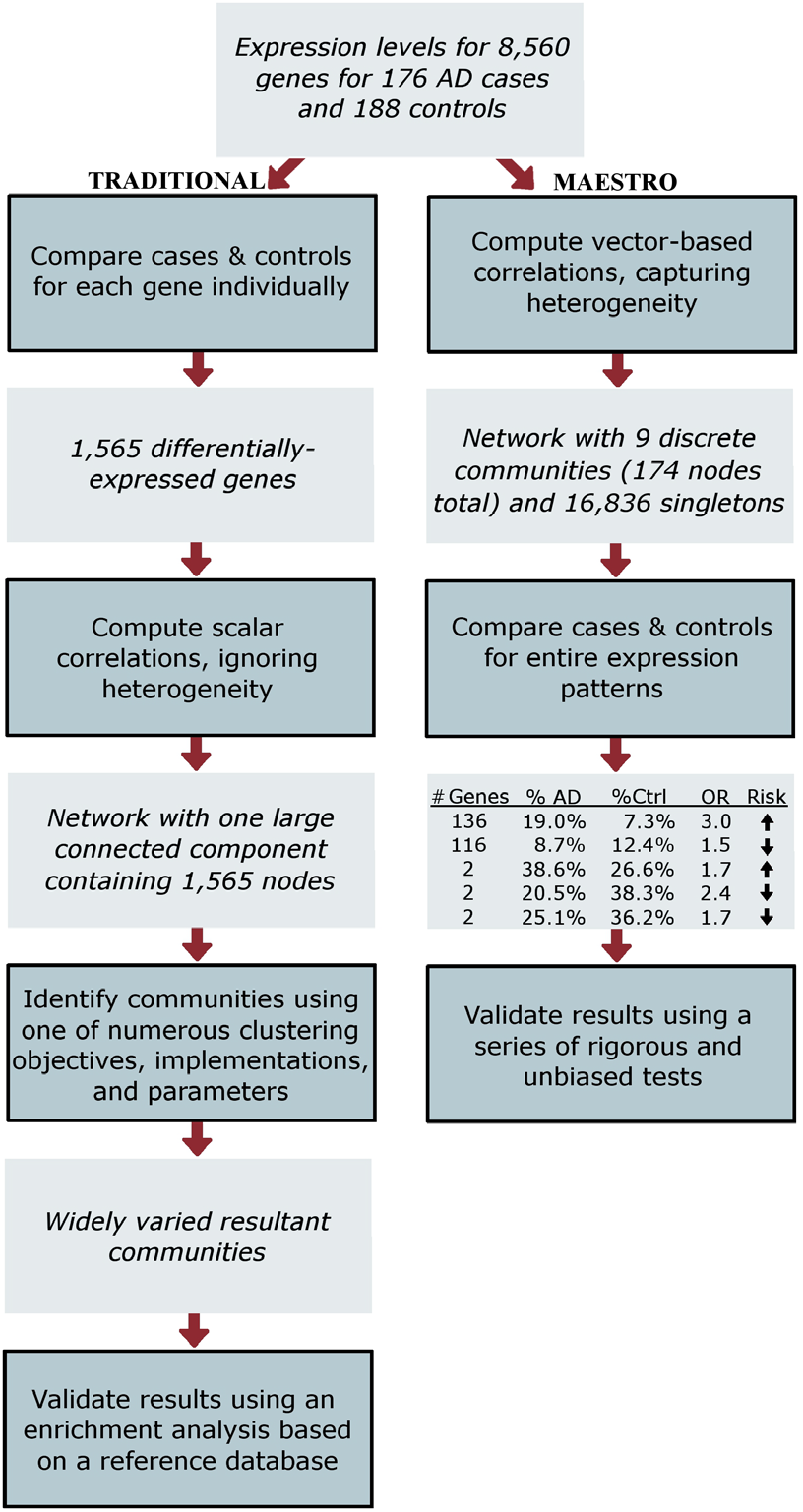
Overview of traditional co-expression and Maestro analyses with highlights of the AD results.

First, the elimination of genes that failed to be detected as differentially expressed led to the loss of 22% of the nodes that appeared in modules in the Maestro network for AD. In other words, many of the genes that appeared correlated by Duo are not differentially expressed when considered in isolation, demonstrating the importance of retaining all genes until entire modules of truly co-expressed genes are identified. Furthermore, many clustering algorithms can be used to partition a co-expression network into modules and we observed that even the use of three highly similar methods, all aimed at optimizing the modularity function^19^, produced diverse results. It is not clear which clustering method is more biologically sound, yet the co-expression network must be partitioned in order to obtain any meaningful information. Maestro is not faced with this dilemma as the modules arise naturally separated in the network. Finally, Maestro is about four times faster than the traditional co-expression network method.

## Discussion

Maestro is similar to our previously introduced BlocBuster^20,21^ method (www.blocbuster.org). BlocBuster is used to analyze single nucleotide polymorphism (SNP) data for complex traits exhibiting heterogeneity. BlocBuster’s unique features have enabled us to identify genetic patterns that have been previously overlooked, and revealed combinatorial genetic patterns with strong associations with hypertensive heart disease^21^ and psoriasis^20^, as well as a coadaptation pattern between vitamin D receptor and skin color genes^22^. Furthermore, analysis of HapMap^23^ data yielded the discovery of remarkable ‘yin-yang’ sequences^24^. Notably, the HapMap data are arguably the most extensively studied genome-wide data in existence, yet exhaustive studies using conventional methods failed to detect this genetic anomaly, demonstrating BlocBuster’s distinctive power for the discovery of combinatorial patterns hidden in genetic data. Maestro adapts several of BlocBuster’s advantageous properties for continuous-valued data: a vector-base correlation metric to capture heterogeneity, an enlarged network scaffold to reduce false-positive signals induced by the transitivity assumption, and a computationally efficient implementation.

Maestro is fundamentally different from traditional co-expression network approaches (Figure 6) and produces distinctive results for several reasons. First, the traditional method selects genes that are differentially expressed between cases and controls in the first step, while Maestro does not compare cases and controls until the last step, where entire modules are statistically tested for their discriminative power for separating cases from controls. Consequently, these two approaches address different biological objectives. The traditional approach limits the investigation to the impact of the trait status on the pattern of gene expression. In contrast, Maestro examines the impact of the pattern of gene expression on trait status, which is more valuable and practical from an epidemiological point of view in terms of diagnostic and risk prediction potential. Furthermore, the loss of genes that are not detected as differentially expressed when examined in isolation may reduce power. A differential-expression signal may be masked when the gene is also involved in unrelated processes. However, such a gene may add discriminative power when examined as a part of an entire module of genes that are synergistically expressing to manifest a trait. For example, 22% of the unique genes in the Maestro modules were discarded in the first step of the traditional approach, reflecting the fact that Maestro can detect genes involved in meaningful interactions that don't have significant marginal effects between cases and controls, while the traditional approach cannot detect such genes. Moreover, the large number of DE trials often results in the identification of many genes: 1,565 for these AD data. With such a large number of candidate genes, it is difficult to gain much biological insight into the complex trait system. In contrast, Maestro identified ten-fold fewer genes being associated with AD trait status, and this much smaller number of candidate genes was highly enriched for genes that had been associated or implicated with AD in previous studies not involving gene expression, as will be discussed later. Furthermore, the traditional analysis also creates a large number of edges that increase the density of the network and contribute to the necessity of employing clustering methods to partition the resultant component. Importantly, this network is prone to hosting dual-expressed nodes (Figure 2), in which sets of genes from different biological pathways are merged due to the assumption of transitivity. The resultant network must be subdivided and it is unclear which of the multitude of diverse options is the most biologically relevant. In contrast, Maestro spontaneously produces networks with strong community structure.

The eminent advantage of this *in silico* discovery of patterns embedded in expression data is the generation of hypotheses that are worthy of further exploration. If the 136 genes in the AD pattern were expressed independently of each other, there is a virtually zero probability (≤1.07×10^−48^) that they would exhibit simultaneous high expression. The manifestation of this large-scale pattern in the human brain (Figure 5 c & d) indicates these expression levels are not independent, and contrarily exhibit a tendency towards lockstep expressions. The appearance of the inverse of this pattern further cross validates this coordination of genetic activities. Notably, this pattern is highly associated with AD, exhibiting an odds ratio of 3.0 and Bonferroni-corrected p-value < 2.3×10^−306^, and has been extensively validated.

AD is a neurodegenerative disease in which the brain accumulates extracellular amyloid plaques and intraneuronal neurofibrillary tangles, and hippocampal and cortical neurons die. Prompted by its association with early-onset familial AD, amyloid-β (Aβ) has been a central research focus for decades^25,26^. The neurotoxicity induced by Aβ is poorly understood and various species may exert their deleterious effects both within neurons and within the interstitial space of the human brain. Another perplexing aspect of AD is its exceptionally long progression. It tends to have a duration of about ten years from diagnosis to death, along with possibly an additional 25 years of progression prior to diagnosis^125^.

Table 1 lists the 136 genes in the largest AD module identified by Maestro and Supplementary Spreadsheet 6 lists these genes as well as their locations, aliases, and NCBI descriptions. Interestingly, 63 of these 136 genes have functions that appear relevant for AD disease progression, including pro-survival mechanisms. We discuss these genes next.

**Table 1.**
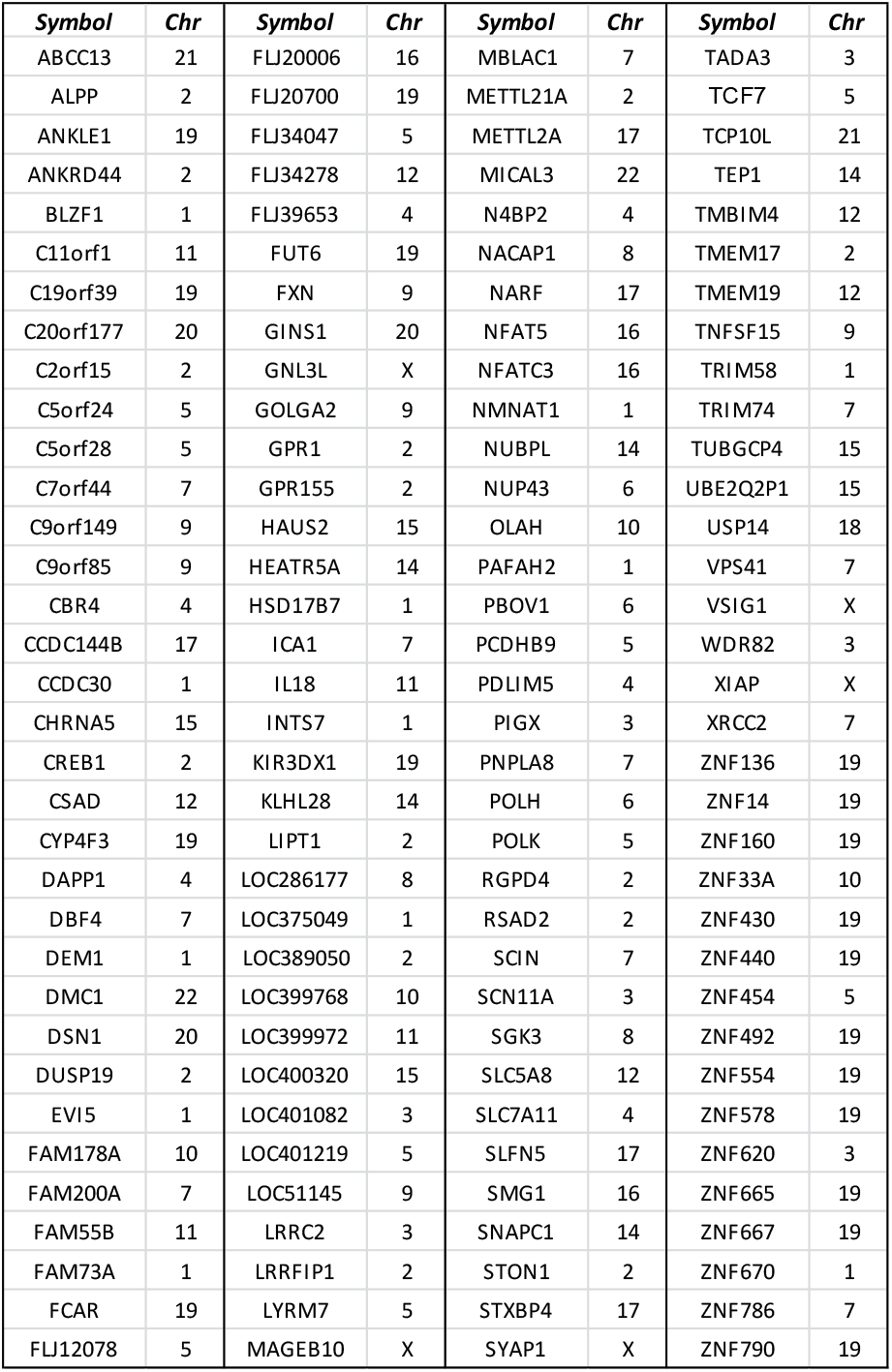
List of 136 genes in the largest module identified for AD. These genes are listed in Supplementary Spreadsheet S6, along with aliases, descriptions, and genomic locations.

Of particular interest is the transcription factor *CREB1* (aka *CREB*), the cAMP-response element binding protein, along with two G-protein coupled receptors (*GPR1* and *GPR155*) which may be involved in the CREB signaling pathway, due to the associations of CREB with AD^27–31^. Mutations in the presenilin genes lead to early-onset familial AD, and it has been demonstrated *in vitro* and *in vivo* that expression of these mutant genes leads to constitutive CREB phosphorylation^28^. While CREB plays key roles in long-term memory formation^32^ and neuronal survival^33^, constitutive CREB activation has been demonstrated to lead to memory deficits^34^, cognitive decline^35^, and neurodegeneration^36^. It has been suggested that constitutive CREB phosphorylation may be involved in AD pathogenesis^28^.

It has been shown that glucocorticoid-induced phosphorylation of CREB is dependent upon serine/threonine-protein kinase, *SGK*^37^, and *SGK3* (aka *CISK*) is also a member of this module. Conversely, CREB has been implicated in the regulation of *SGK3* during apoptosis^38^, and CREB has also been suggested to regulate the following module members: *TNFSF15* (aka *VEGI*), *HSD17B7*, *POLK*, *IL-18*, *DUSP19*, *RSAD2*, *ICAI* (aka *ICA69*), *KLHL28* (aka *BTBD5*), *SLC5A8*, and *ZNF667* (aka *MIPU1*).

DNA damage has been widely noted in AD brains with observations as early as 1990^39^, and a number of genes in the 136-gene module are involved with DNA damage response. Genes encoding two of the Y-family DNA polymerases, *POLH* (aka *XPV* and *RAD30a*) and *POLK*, are members of this module. Y-family polymerases are specialized for replicating past DNA lesions, but are highly regulated, as they have low fidelity for undamaged DNA^40^. It should be noted that there are only four known Y-family polymerases, and two are members of the 136-gene module.

Module members *XRCC2*, *ANKLE1* (aka *LEM3*), *GINS1* (aka *PSF1*), *NMNAT1*, *DBF4*, *SMG1*, and *DEM1* (aka *EXO5*) have also been associated with DNA damage^41–45^. *XRCC2* encodes a member of the RAD51 heterotetramer, and its mutation leads to hypersensitivity to double strand breaks^44,46^. *ANKLE1* is a conserved endonuclease that is capable of inducing DNA damage response and its loss in *C. elegans* results with hypersensitivity to DNA damage^42,47^. *GINS1* also induces DNA damage response^48^. NMNAT1 binds to PARP-1, thereby amplifying poly-ADP-ribosylation in response to single-strand breaks^45^. *DBF4* activates Cdc7 kinase during DNA repair^49^. *SMG1*, a member of the phosphatidylinositol 3-kinase (PI3K)-related kinase family, is involved in p53-mediated response to genotoxic stress^50^. *DEM1* is an exonuclease that appears to be essential for genomic stability in yeast mitochondria^41^.

The DNA damage response involves histone modifications, including methylation of histone H3 lysine 9 (H3K9) for efficient repairs^51,52^. A methyltransferase, *SUV39H2*, which is important for genomic stability and preferential to H3K9 methylation^53–55^, is a member of the 136-gene module. Some additional module genes that may potentially be relevant are *WDR82*, an H3K4 methyltransferase; *C11orf1*, which binds to the H3K9 methyltransferase *SETDB1*; two methyltransferase-like genes, *METTL2A* and *METTL21A*^56,57^; and *TADA3* (aka *ADA3*), which is a transcriptional activator adapter that is part of the p300/CREB binding protein (PCAF) histone acetylase complex and it is also critical for full activation of p53^58^. Interestingly, PCAF knockout mice exhibit a resistance to Aβ toxicity^59^.

Over the last few years several key impacts of intraneuronal accumulations of Aβ on AD progression have been illuminated^60–62^. While the sources of these accumulations are not well understood, several mechanisms for Aβ uptake from extracellular space have been explored, including endocytosis of Aβ bound to nicotinic acetylcholine receptor (nAChR)^63–65^. It has been shown that Aβ binds to the α7 subunit of nAChR with picomolar affinity, the complex is rapidly internalized, and elimination of this path blocks Aβ internalization for *in vitro* studies^65^. Using a transgenic mouse model, it was shown that deletion of the α7 nAChR gene protected the brain from the cognitive deficiencies associated with AD^64^. Furthermore, nicotine stimulation of α4β2 nAChR in cultured rat cortical neurons exhibited a protection from Aβ toxicity^66^ and Aβ blocked α4β2, α2β2, and α4β2α5 nAChRs in *Xenopus* oocytes^67^. In general, cholinergic signaling has historically been of central interest in the progression of AD and a number of AD drugs are acetylcholinesterase inhibitors^68^; plus at least five additional AD drugs target nAChRs^69^.

A potentially relevant gene in the 136-gene module is *CHRNA5*, which encodes α5 nAChR. *CHRNA5* is associated with lung cancer as well as nicotine addiction^70,71^. In the brain, it is most commonly found in α4β2α5 nAChR and the presence of the α5 subunit alters cellular trafficking^72^, increases the receptor’s function without impacting it’s expression^73^, and affects α4β2_*_ nAChR interactions with synaptic scaffolding proteins^74^. Furthermore, α5 inclusion also increases the rate of desensitization and increases Ca^2+^ permeability of α4β2^*^ nAChR^72,75^. Reductions in the numbers of nAChRs on cell surfaces have been associated with AD^69^, yet *CHRNA5* exhibits high expression in the risk pattern. Due to the rapid turnover rate of the nAChR/Aβ complex^65^, high expression of nAChR genes with low surface density of these receptors could be explained by a high rate of Aβ internalization.

A number of studies have indicated that significant mitochondrial dysfunction occurs early in the progression of AD, and Aβ accumulation within mitochondria has been implicated in this dysfunction^61,76–78^. It has been shown that Aβ is transported into the mitochondria via the TOM machinery^76^ where it has the potential to contribute to increased oxidative stress, disruption of cellular Ca^2+^ homeostasis, synaptic degeneration, and initiation of apoptotic pathways^61,77–83^. Dysfunctional mitochondria in AD neurons may activate the caspase-9/-3 apoptotic pathway^84,85^. Furthermore, *in vitro* microinjections of intracellular Aβ in human neurons result in rapid apoptosis via the p53-Bax pathway^86,87^. However, the long duration of AD progression suggests that pro-survival mechanisms must interfere with these apoptotic pathways and commence so-called abortosis^88,89^.

Additional potentially relevant genes in the 136-gene module include *TMBIM4*, *XIAP*, *TNFSF15*, *TEP1*, *NFAT5*, and *NFATC3*. *TMBIM4* (aka *GAAP*) encodes a six-transmembrane protein that inhibits apoptosis stimulated by Bax, Fas, and TNF^90,91^. High expression of *TMBIM4* has also been shown to alter Ca^2+^ fluxes and lower the effectiveness of InsP_3_^92^. *XIAP* encodes an X-linked inhibitor of apoptosis protein (IAP) that blocks caspases 3, 7, and 9^93^. *XIAP* may be the only true caspase inhibitor in the IAP family^94^, and it is a critical discriminator of Fas types I and II apoptosis signaling^95^. Furthermore, *XIAP* phosphorylation by protein kinase C may be a prosurvival mechanism for AD^96^. *TNFSF15* (aka *VEGI*) encodes a cytokine of the tumor necrosis factor (TNF) ligand family. TNFSF15 is the only known ligand for TNFRSF25 (aka Death Receptor 3) and the complex can lead to either apoptosis or activation of the NFκB/MAPK pathway^97^. *TEP1* encodes a telomerase that compensates for the shortening of telomeres due to cell divisions^98^. In general, telomerases are not expressed beyond postnatal development, with the exception of cancerous or stem cells^99^. It has been shown that telomerase expression is capable of interrupting apoptosis induced by Aβ prior to mitochondrial dysfunction^100^, and also that *TEP1* interacts directly with p53^101^. (Interestingly, longer telomeres have been associated with the AD risk allele, *APOEε*4, and the coupling of this allele with longer telomeres predicted diminished episodic memory^102^.) Nuclear factor of activated T-cells (NFAT) has been shown to regulate apoptosis via both NFAT/Fas^103^ and Ca^2+^/calcineurin/NFAT signaling by p53^104^, and *NFAT5* and *NFATC3* are module members.

Recent discoveries have shed light on the activities of Aβ after translocation into the mitochondria. ABAD, the Type 10 member of the 17-beta-hydroxysteroid dehydrogenase (17β-HSD) family, is normally bound by nicotinamide adenine dinucleotide (NAD) in neuronal mitochondria and catalyzes reversible oxidation and/or reduction of alcohol groups^105^. However, Aβ competes with NAD for ABAD binding^61,105^. Multiple studies have found that inhibiting the Aβ-ABAD interaction reduces AD impairment^61,78,106^. In addition to its role in DNA repair (see above), NMNAT1 is the master enzyme of NAD synthesis. Surprisingly, NMNAT1 has also been shown to have chaperone activities with neuronal protection in response to stress^107^. Notably, it has been demonstrated to promote clearance of hyperphosphorylated Tau in *Drosophila*^108^. Hyperphosphorylated Tau is the primary component of the neurofibrillary tangles that are characteristic of late-stage AD. *NMNAT1* is a member of the 136-gene module, along with Type 7 member of 17β-HSD, *HSD17B7* (aka *PRAP*); as well as *CBR4*, which couples with Type 8 member of 17β-HSD in catalyzing the second step of mitochondrial fatty acid synthesis^109^.

There are a number of additional interesting genes in this module. *SLC7A11* (aka *xCT*) encodes a member of the Xc(-) glutamate transporter system, which increases microglial-associated cytotoxicity of Aβ, while also masking potential neuroprotection via microglial APOE^110^. *GOLGA2* has been suggested to play a key role in the fragmentation of the Golgi due to AD deregulation^111^. *PDLIM5* is a homolog of *AD7c-NTP*, which encodes a neuronal thread protein that has been utilized as an AD biomarker^112,113^. Genes associated with inflammation and/or immune response include: *CYP4F3*^114^, *FCAR*^115^, *SCN11A*^116^, *TNFSF15*^117^, *KIR3DX1*^118^, *NFAT5*^119^, *NFATC3*^120^, and *IL18*^121^. *IL18* has also been associated with AD in the Han Chinese population^122^. The module also includes the following synapse-related genes: *SCIN*^123^, *STON1*^124^, *STXBP4*^125^, *USP14*^126^, and *SYAP1*^127^. Finally, the module possesses 17 genes with zinc finger domains (for relative comparisons, 1.16% of the gene symbols in the entire 8,560 gene set and 4.41% in the 136-gene pattern began with ‘ZNF’, suggesting an enrichment.)

Taken together, the membership of this module includes 63 genes that immediately appear to be of interest for the study of AD. Many of these are pro-survival genes, suggesting cells fighting for life. This hypothesis is in line with the extremely slow progression of AD, as it tends to have a duration of more than 30 years^128^. The synchronization of expressions within the module of 136 genes represents synchronized coordination of genetic activities in the human brain and establishes Maestro’s adeptness for exposing combinatorial co-expression patterns embedded within genome-wide data. This pattern also exhibits a highly significant association with AD, paving the way for future breakthroughs in our understanding of the progression of this devastating disease.

In conclusion, it should be noted that Maestro can be applied to the study of any complex trait as well as diverse domains in which heterogeneity is present.

## Experimental Procedures

### Material

The sample of AD cases and normal controls, as well as the companion gene expression data, have been described in a previous study that we were involved with^12^ and are summarized in the SI. Expression data are available from the Laboratory of Functional Neurogenomics at http://labs.med.miami.edu/myers/LFuN/data%20ajhg.html.

### Maestro

Following is a brief description of the methods used for Maestro.

#### Duo

Duo is a simple metric that efficiently computes correlations for every pair of genes, not just those that are differentially expressed. We considered four types of interesting relationships between a pair of genes: both genes have high expression (HH), both have low expression (LL), or the genes are anti-correlated (HL or LH). The computation of each of the Duo values begins with the percentage of individuals exhibiting one of the four relationships, denoted as *R_ij_*[*t*], where *t* is the relationship type and *i* and *j* are the gene pair. The goal of a correlation measure is to quantify the departure from independence exhibited by the values of two variables. For this reason, the *R_ij_*[*t*] values are multiplied by two frequency factors (one for each gene), thereby accounting for the effect of the frequencies upon the probability of observing the relationship by random chance. The current implementation of Maestro uses a simple frequency factor, *ff_it_*, for this purpose. For gene *i*, this factor was set to 1 − *f_it_*/*q*, where *f*_*it*_ is the frequency of the relevant expression direction (e.g. high expression) for relationship *t* for the group of individuals and *q* is a weighting factor, which was set to 1.5 in this prototype implementation. We scaled the product by a factor of four in order to have the values approach a range of zero to one, resulting with the following formula: *DUO_ij_*[*t*]= 4 *R_ij_*[*t*] *ff*_*it*_*ff*_*jt*_ for each pair of genes, *i* and *j*, and their relationship *t*.

#### Network construction

In the network, each gene was represented by a pair of nodes, representing high and low expression, respectively. Duo was used to measure correlations between each of a pair of nodes. An edge was placed between the corresponding nodes in the network for the 1,000 highest correlation values. Module memberships were identified using a breadth-first search.

#### Carriers algorithm

We developed a Carriers algorithm for computing the variation of expression of entire patterns between cases and controls. For each pattern, Carriers tallied the numbers of individuals that exhibited the entire expression pattern and compared the two groups using odds ratio and corrected p-value based on the G-test.

*Statistical analyses* are described in the SI.

### Validation Trials

#### Permutation trials

We used permutation trials to test the probability that a pair of genes would be labeled as correlated by random chance. In these trials the expression values of the individuals were randomly shuffled for each gene. Consequently, each gene had its original mean, deviation, and other statistical properties; the only change was that the inherent “correlations” between pairs of genes had been removed by rearranging the expression values across individuals. After each randomization, we computed the Duo correlation values for every pair of genes. Given this randomization scheme, we would expect that no significant correlations should remain in the data.

#### Replication of results

We randomly split the data into two sets, each of which had data for 88 cases and 94 controls. We combined the cases and controls data for the first half and built a 1000-edge network using Duo and identified the connected modules. The identified patterns were tested on the other half of the individuals.

#### Variation of network density

We varied the number of edges in the networks to test the robustness of the modules to our somewhat arbitrary choice of 1,000 edges. We ran four additional trials, in which we reduced the number of edges to 500 and then increased it to 5,000, 10,000, and 15,000.

#### Segregation of data with known heterogeneity

We demonstrated Maestro’s ability to segregate heterogeneity by manually segregating known heterogeneity and testing the subsets. We separated AD cases from controls and built two networks, one for each group of individuals.

#### Visual inspection of expression patterns

Carriers tallies the individuals that possess the entire expression pattern, but does not capture individuals that possess most of the pattern. It is possible that partial patterns might be more prominent in one group than the other, potentially weakening the significance of the association. In order to detect these partial patterns, we plotted all of the expression values for the genes within the pattern of interest and visually inspected the results.

## Acknowledgments

We gratefully acknowledge Cynthia C. Vigueira, Patrick Vigueira, Alison Goate, and Oscar Harari for valuable suggestions; Amanda Myers for supplying data; and funding from the National Institutes of Health (1RF1AG053303-01, R01GM08641201, P50-GM65509, RC1AR05868102, R01HL09102801), National Science Foundation (DBI-0743797), Knight Alzheimer’s Disease Research Center (ADRC) Pilot Grant, and the Alzheimer’s Association.

