## Supplementary File for "Synchronized genetic activities in Alzheimer’s brains revealed by heterogeneity-capturing network analysis"

### SUPPLEMENTARY TEXT AND FIGURES

***Six Excel spreadsheets available in separate files.***

**Supplementary Table S1.** Description of the Maestro 1k network built from 176 AD cases and 188 controls. There were 16,836 singletons (nodes with no incident edges) and five doubletons (a pair of nodes joined by a single edge). The number of nodes and edges for each of the other modules are shown. The average density of the modules was 0.70, where density is defined as the number of edges divided by the number of possible edges.

| Module | # Nodes | # Edges |
|---|---|---|
| 1 | 136 | 583 |
| 2 | 116 | 371 |
| 3 | 19 | 38 |
| 4 | 3 | 3 |

**Supplementary Table S2.** Summary of results for the two largest modules identified in the discovery dataset and tested on both the discovery and replication data. Number of nodes and edges in each module, expression direction, number of individuals possessing pattern, percent of cases and controls possessing pattern, odds ratio, and Bonferroni-corrected p-values are shown.

| # Nodes | # Edges | Expression Direction | ***Discovery Data*** | | | | | ***Replication Data*** | | | | |
|---|---|---|---|---|---|---|---|---|---|---|---|---|
| | | | # Indiv | % Cases | % Ctrls | OR | p-value | # Indiv | % Cases | % Ctrls | OR | p-value |
| 113 | 405 | High | 15 | 13.1 | 4.7 | 3.1 | $<2.3x10^{-306}$ | 23 | 20.0 | 7.3 | 3.2 | $<2.3x10^{-306}$ |
| 113 | 461 | Low | 20 | 10.5 | 12.0 | 1.2 | $<2.3x10^{-306}$ | 18 | 5.7 | 14.3 | 2.8 | $<2.3x10^{-306}$ |

**Supplementary Table S3**. Variation of number of edges for Maestro networks. Summary of networks with 500, 1k, 5k, 10k, and 15k edges built from AD cases and controls data. The number of nodes with no incident edges (singletons), number of modules that are comprised of a single edge connecting two nodes (doubletons), and number of modules with at least three nodes are listed. For each of the three modules with the most edges in each network, the number of nodes and edges in the modules as well as the number of cases and controls possessing the entire corresponding expression patterns, resultant odds ratios, and Bonferroni-corrected p-values are shown.

| | Number of Edges in Network | 500 | 1k | 5k | 10k | 15k |
|---|---|---|---|---|---|---|
| **Numbers of Modules** | ***# Singletons*** | 16,908 | 16,836 | 16,642 | 16,531 | 16,419 |
| | ***# Doubletons*** | 7 | 5 | 10 | 11 | 19 |
| | ***# 3 or More Nodes*** | 5 | 4 | 6 | 9 | 12 |
| **Largest Module** | ***# Nodes*** | 105 | 136 | 212 | 242 | 268 |
| | ***# Edges*** | 319 | 583 | 2,791 | 5,420 | 7,958 |
| | ***# Cases (%)*** | 33 (19.0%) | 33 (19.0%) | 19 (11.7%) | 17 (11.2%) | 15 (10.7%) |
| | ***# Controls (%)*** | 16 (8.7%) | 13 (7.3%) | 6 (3.9%) | 6 (4.6%) | 5 (4.2%) |
| | ***Odds Ratio*** | **2.4** | **3.0** | **3.3** | **2.6** | **2.7** |
| | ***p-value*** | **< 2.3E-306** | **< 2.3E-306** | **< 2.3E-306** | **< 2.3E-306** | **< 2.3E-306** |
| **2nd Largest Module** | ***# Nodes*** | 74 | 116 | 185 | 220 | 341 |
| | ***# Edges*** | 153 | 371 | 2,011 | 4,123 | 6,965 |
| | ***# Cases (%)*** | 18 (10.3%) | 15 (8.7%) | 10 (5.9%) | 6 (3.6%) | 1 (1.2%) |
| | ***# Controls (%)*** | 26 (14.4%) | 22 (12.4%) | 14 (8.0%) | 12 (7.2%) | 2 (1.5%) |
| | ***Odds Ratio*** | **1.5** | **1.5** | **1.4** | **2.1** | **1.3** |
| | ***p-value*** | **< 2.3E-306** | **< 2.3E-306** | **< 2.3E-306** | **< 2.3E-306** | **< 2.3E-306** |
| **3rd Largest Module** | ***# Nodes*** | 12 | 19 | 50 | 75 | 17 |
| | ***# Edges*** | 15 | 38 | 180 | 420 | 27 |
| | ***# Cases (%)*** | 37 (24.0%) | 30 (22.4%) | 16 (13.7%) | 10 (9.9%) | 31 (19.3%) |
| | ***# Controls (%)*** | 51 (28.2%) | 38 (21.5%) | 21 (12.5%) | 14 (8.9%) | 20 (10.8%) |
| | ***Odds Ratio*** | **1.2** | **1.1** | **1.1** | **1.1** | **2.0** |
| | ***p-value*** | **< 2.3E-306** | **< 2.3E-306** | **< 2.3E-306** | **< 2.3E-306** | **< 2.3E-306** |

**Supplementary Table S4.** Description of the Maestro network built using AD cases data only (176 individuals). There were 16,952 singletons (nodes with no incident edges). The number of nodes and edges for each of the other modules are shown. There were a total of 1,002 edges due to ties in the Duo values. The average density of the modules was 0.69.

| Module | # Nodes | # Edges |
|---|---|---|
| 1 | 164 | 1000 |
| 2 | 2 | 1 |
| 3 | 2 | 1 |

**Supplementary Table S5.** Description of the Maestro network built using AD controls data only (188 individuals). There were 16,953 singletons (nodes with no incident edges). The number of nodes and edges for each of the other modules are shown. There were a total of 1,001 edges due to a tie in the Duo values. The average density of the modules was 0.69.

| Module | # Nodes | # Edges |
|---|---|---|
| 1 | 160 | 997 |
| 2 | 3 | 2 |
| 3 | 2 | 1 |
| 4 | 2 | 1 |

#### Supplementary Experimental Procedures

**Material.** The sample of AD cases and normal controls, as well as the companion gene expression data, have been described in a previous study that we were involved with[1]. Briefly, subjects were self-defined European descent, at least 65 years old at time of death, and neuropathological diagnoses were based on standard NACC protocols. Illumina Human Refseq-8 Expression BeadChips were employed with standard protocols. Illumina BeadStudio software was utilized and rank-invariant-normalized expression values were $\log_{10}$ transformed, with missing data encoded as missing (not zero). Only transcripts that were detected in at least 90% of the cases or controls were retained in order to enrich for primary effects. Data were corrected for gender, age at death, cortical region, APOE status, day of expression hybridization, institute source of sample, postmortem interval, and transcription detection rate.

**Alternative co-expression network construction**. For comparative purposes, we analyzed the AD data using a well-established co-expression network method, HQcut. Specifically, the method and parameters described in our previous paper on AD[2] were used. Briefly, we first identified differentially expressed (DE) genes using SAM[3]. Pearson's correlation coefficient (PCC) was computed for every pair of these differentially-expressed genes. An edge was placed between two nodes if their expression profiles satisfied at least one of the two following criteria: (1) the absolute value of their Pearson's correlation coefficient (|PCC|) was higher than 0.3 and, for at least one of the two genes, the |PCC| value was amongst the highest three values that it had with any other gene in the network, or (2) their |PCC| was higher than 0.8 and it was amongst the highest 50 |PCC| values.

Traditional co-expression networks are generally comprised of a single large module connecting all of the DE genes. Clustering methods are utilized to partition this large module into smaller modules with strong interconnectivities. There are a number of clustering strategies that optimize various objectives, such as *k*-means, hierarchical clustering, and the modularity function $Q$[4]. It is not clear which objective is the biologically most meaningful for partitioning co-expression networks. However, *k*-means and hierarchical clustering enforce sphericity, in which they exhibit a bias toward hyperspherical clusters, and there is no reason to expect these biological clusters to possess this property. The modularity function $Q$ does not enforce this assumption and the objective is aimed at identifying module structures in networks, in which there are many intra-cluster edges and few inter-cluster edges[5]. Furthermore, modularity appears to arise naturally in gene expression networks[6] and it has previously been employed for the partitioning of such networks[7,8].

We applied three different clustering methods that all strive to optimize the modularity function $Q$ via various spectral graph partitioning computational approaches: Newman's implementation[5], QCUT[9], and HQCUT[9]. In order to quantify the variations between cluster memberships, we computed the Jaccard Index for all pairs of clusters produced by each of the methods. The Jaccard Index is defined as follows. Let $S_j = \{(u,v) : u,v \in c_j\}$ be the set of all pairs of vertices that appear together in any cluster, $c_j$. Let $N_i$ equal the size of $S_j \cap S_k$, and $N_u$ equal the size of $S_j \cup S_k$, for clusters $j$ and $k$. The Jaccard Index is calculated by $JI = N_i / N_u$ and ranges from zero, for two clusters with no nodes in common, to one, for two identical clusters. Comparisons of results are discussed below.

**Statistical analyses.** Since the network was blindly built using cases and controls data, it could be expected that some patterns are common at the population level, with no association to the phenotype. However, because Duo accommodates genetic heterogeneity, it might also be expected that some of these patterns exhibit associations with the phenotype status, which is one type of heterogeneity. Two metrics were used to evaluate the significance of these patterns: odds ratio and G-test. Specifically, we used the following formula for the odds ratio: $OR = p(1\text{-}q) / q(1\text{-}p)$, where $p$ and $q$ equal the percentages of cases and controls possessing the entire pattern, with $p \geq q$. (This formulation yields values greater than one for both risk and protective patterns for ease of comparisons.) Individuals that have missing expression values for any of the genes in the given pattern were not included in the computations.

G-test is a maximum likelihood statistical significance test, that is recommended for this type of study and appears to approximate the $\chi^2$ distribution more accurately than Pearson's $X^2$ metric[10]. We used the following formula for the G-test goodness of fit: $G = 2\sum_i O_i \cdot \ln(O_i / E_i)$ where $O_i$ and $E_i$ equal the observed and expected number of individuals possessing the entire pattern in subset $i$. The summation is over the two subsets, cases and controls, and 'ln' denotes the natural log. The expected numbers of individuals were computed by multiplying the frequencies of expressions of the relevant genes for each group. The p-value of significance corresponding to the G-test score, with Bonferroni correction for multiple testing, was used for each result.

#### Supplementary Text

**Spurious correlations due to small sample size.** It should be noted that spurious edges can arise due to small sample size. In our preliminary trials using *Saccharomyces cerevisiae* data consisting of gene expression values for 6,228 genes for 44 samples (unpublished data), we identified an average of 6,304 edges with Duo values ≥ 0.8 per permutation trial. These results were highly consistent, with a maximum of 6,636 edges and minimum of 6,064 edges over 1,000 trials. Duo had produced more edges for the original *S. cerevisiae* data, but the permutation trials indicated that thousands of these may be due to random chance.

We ran 1,000 permutation trials for the AD dataset and there were no correlations with Duo values greater than 0.524. We suspected the small sample size of the *S. cerevisiae* dataset was the primary cause for the appearance of random edges in the earlier permutation trials, and the 364 individuals in the AD dataset were adequate to eliminate these false positive signals. To test this hypothesis we randomly extracted 44 samples from the original AD data and conducted the permutation validation. This trial had very similar results to our findings for *S. cerevisiae*, suggesting small sample size can yield false positive edges. The minimum number of samples needed likely varies for different datasets. For these reasons, permutation tests should be run whenever correlations are being computed to ensure that false positive edges are not produced.

**Comparison with co-expression network method.** We analyzed the data using a well-established co-expression network method[8] (see SI) and comparisons between Maestro and the traditional approach are summarized in Figure 6 in the manuscript. The resultant AD co-expression network had 1,565 nodes and 15,629 edges. We discuss several key differences between the results for the co-expression method and Maestro next.

First, we observed that despite the fact that 1,565 of the 8,560 AD genes were identified as differentially expressed by SAM[3] and retained in the co-expression network, 22% of the 109 unique genes that had incident edges in the Duo network were eliminated by SAM as they did not exhibit significant differential expression when considered in isolation. Elimination of genes that are not differentially expressed is a common first step of co-expression methods. However, as shown here, these eliminations can result with the loss of genes that may be of value for the research effort.

Second, because PCC provides a single scalar that disregards genetic heterogeneity, it has reduced discriminatory power and may introduce false positive edges, as shown in the results for the permutation trials, as described above. Moreover, the selection of a biased subset of genes, those that are differentially expressed, prior to building the network increases the probability of inter-correlations due to their shared differential expressions.

Third, the co-expression method requires the application of a clustering method as the constructed network typically possesses a single connected component, in which every node is connected. We applied three different clustering methods that all strive to optimize the modularity function *Q*[4] via various spectral graph partitioning computational approaches: Newman's implementation[5], QCUT[9], and HQCUT[9]. The resultant clusters that were identified vary widely for the three methods, and they produced 14, 8, and 103 clusters for Newman's implementation, QCUT, and HQCUT, respectively. In order to quantify the variations between cluster memberships, we computed the Jaccard Index for all pairs of clusters produced (e.g. Newman's vs. QCUT). For each cluster in a given result we computed the highest Jaccard Index over all the clusters found by the other method. The cluster memberships varied substantially as the averages of these maximum Jaccard Indices were 0.353, 0.087, 0.064 for comparisons between each pair of the three methods. QCUT and HQCUT, which were developed together, had the highest of these averages (0.353) and Newman's results had little in common with either of the other two results.

In addition to spectral clustering, a number of other computational methods have been employed to optimize modularity $Q$ such as a greedy divisive algorithm[4], a divisive extremal optimization approach[11], simulated annealing approaches[12], and agglomerative hierarchical clustering algorithm[13]. Overall, there are many techniques available for clustering the nodes in the co-exp network, and they yield quite diverse results.

In summary, the elimination of genes that failed to be detected as differentially expressed led to the loss of 22% of the nodes that appeared in modules in the Maestro network for AD. In other words, many of the genes that appeared correlated by Duo are not differentially expressed when considered in isolation, demonstrating the importance of retaining all genes until entire modules of co-expressed genes are identified. Furthermore, many clustering algorithms can be used to partition a co-expression network into modules and we observed that even the use of three highly similar methods produced diverse results. It is not clear which clustering method is more biologically sound, yet the co-expression network must be partitioned in order to obtain any meaningful information. Maestro is not faced with this dilemma as the modules arise naturally separated in the network.
